# Identification of transcriptional regulators using a combined disease module identification and prize-collecting Steiner tree approach

**DOI:** 10.1101/2024.02.05.577574

**Authors:** Gihanna Galindez, Ben Anthony Lopez, David B. Blumenthal, Tim Kacprowski

## Abstract

Transcription factors play important roles in maintaining normal biological function, and their dysregulation can lead to the development of diseases. Identifying candidate transcription factors involved in disease pathogenesis is thus an important task for deriving mechanistic insights from gene expression data. We developed Transcriptional Regulator Identification using Prize-collecting Steiner trees (TRIPS), a workflow for identifying candidate transcriptional regulators from case-control expression data. In the first step, TRIPS combines the results of differential expression analysis with a disease module identification step to retrieve perturbed subnetworks comprising an expanded gene list. TRIPS then solves a prize-collecting Steiner tree problem on a gene regulatory network, thereby identifying candidate transcriptional modules and transcription factors. We compare TRIPS to relevant methods using publicly available disease datasets and show that the proposed workflow can recover known disease-associated transcription factors with high precision. Network perturbation analyses demonstrate the reliability of TRIPS results. We further evaluate TRIPS on Alzheimer’s disease, diabetic kidney disease, and prostate cancer single-cell omics datasets. Overall, TRIPS is a useful approach for prioritizing transcriptional mechanisms for further downstream analyses.

## 1 Introduction

Gene regulation is complex and tightly regulated. Transcriptional regulatory programs are primarily controlled by transcription factors (TFs), specialized proteins that bind to specific DNA sequences to influence the expression patterns of downstream target genes. TFs orchestrate and fine-tune regulatory programs to control fundamental biological processes, including intracellular signaling crucial for determining cell fate, metabolism, and response to environmental stimuli [1–5]. As a result, transcriptional networks are context-specific and thus vary across tissues [6], disease states [7], and time points. Dysregulation in the gene expression programs controlled by TFs can lead to the development of diseases [8, 9]. For instance, TF activity is altered in various cancer types [9]. Given their central roles in maintaining normal cellular functions, TFs are frequent drivers of disease and represent promising therapeutic targets [10, 11]. Consequently, mining candidate TF regulators from omics data has been an active area of research.

The conventional method of investigating the mechanisms behind diseases of interest involves performing differential expression analysis (DEA), which aims to identify genes by looking at significant changes in expression levels between two distinct patient groups. However, simply analyzing gene lists ignores the complex and molecular interactions that occur in the cells and provides a limited picture of the regulatory landscape for a specific biological context [12, 13]. Network-based methods that account for gene-gene interactions provide a natural framework to facilitate a more mechanistic interpretation of experimental results. A common methodological principle of network-based methods is to map gene lists determined from experiments onto template molecular networks, followed by the application of graph-based algorithms that identify subgraphs representing perturbed disease modules. These methods break down complex disease states into distinct functional modules by considering biological signals in aggregate, such that weak signals from individual gene scores are amplified, effectively narrowing down hundreds of genes into more compact subnetworks potentially responsible for the physiological traits of interest. [14, 15].

Along with significant advances in high-throughput omics technologies, various network-based approaches for analyzing experimental data have been developed over the past decade [16, 17]. The accumulating wealth of experimental knowledge has increasingly facilitated the use of tools that leverage prior knowledge derived from curated repositories, often in the form of reference molecular networks (e.g. PPI networks, signaling networks, and gene regulatory networks), to guide the discovery of disease-associated mechanisms [18, 19]. However, results of a recent benchmarking study revealed that most existing tools that utilize template molecular networks fail to effectively use the network structure and instead identify candidate modules primarily based on node degrees [20]. While the tools were originally evaluated in the context of disease module identification, these important findings highlight the general need for developing more reliable methods that effectively use prior knowledge encoded in the network structure to generate predictions.

In this study, we propose Transcriptional Regulator Identification using Prize-collecting Steiner trees (TRIPS), a workflow designed for the task of TF prioritization. We hypothesized that using the prize-collecting Steiner tree (PCST) algorithm as the main component of the method is a suitable approach that can effectively incorporate experimentally derived biological signals to discover a compact, context-specific transcriptional module while effectively utilizing network structure information. In addition, combining protein-protein interactions and gene regulatory interactions in a biologically principled manner can provide additional molecular context. The first step of TRIPS involves disease module identification on a protein-protein interaction (PPI) network. Next, using an augmented gene list from the first step, the PCST problem is solved on a template gene regulatory network (GRN). We evaluate the performance of the proposed workflow on a range of diseases and demonstrate its reliability based on network perturbation analysis. Given its interpretable approach to TF mining, TRIPS can be used to generate mechanistic hypotheses and help shed light into the molecular mechanisms underlying disease that can be prioritized for down-stream analyses.

## 2 Materials and Methods

### 2.1 Dataset collection

#### Evaluation datasets

We analyzed 22 disease-associated RNA-seq datasets with cases and controls across 16 disease phenotypes. Count data of uniformly preprocessed datasets and differentially expressed genes were obtained using the publicly available GREIN resource [21]. Genes were considered differentially expressed at a log2 fold change > 1 and adjusted p-value < 0.05. Detailed information on the datasets used for the analyses are provided in (Supplementary Table S1).

#### Alzheimer’s disease single-cell dataset

DEA results from single-cell RNA-seq (scRNA-seq) from Alzheimer disease (AD) “homeostatic” and “activated” microglial cell populations were obtained from [22]. Genes were regarded as differentially expressed at ln fold change > 0.25 and adjusted p-value < 0.05.

#### Alzheimer’s disease paired single-nucleus RNA-sequencing (snRNA-seq) and single-nucleus ATAC-sequencing (snATAC-seq) dataset

We downloaded cell type-specific DEA results derived from snRNA-seq from microglia isolated from the brains of AD patients and healthy controls. For the same set of samples, chromatin accessibility was profiled using snATAC-seq, and TF motif enrichment of differentially accessible regions (DARs) were determined. The corresponding tables were downloaded from Supplementary Data 1a and 1d of [23]. Genes were regarded as differentially expressed at log fold change > 0.25 and adjusted p-value < 0.01. The DEA results were used as input to TRIPS, PCSF, and MOGAMUN, while voom-transformed raw counts were used as input to CaCTS and RegEnrich.

#### Diabetic kidney disease dataset

We evaluated the different TF prioritization methods using paired single-nucleus RNA-seq and snATAC-seq data in diabetic kidney disease (DKD) profiled across different kidney cell types. We analyzed the following scRNA-seq cell types: *CLDN16*(-) thick ascending limb, *CLDN16*(+) thick ascending limb, endothelial cells, *VCAM1*(+) proximal tubule cells, podocytes, ICA-type A intercalated cells, ICA-type B intercalated cells, principal cells, early distal convoluted tubule, late distal convoluted tubule, ascending thin limb, and parietal epithelial cells. For the same set of samples, the authors profile chromatin accessibility using snATAC-seq and performed TF motif enrichment of DARs. The TFs with significant DAR were used as the ground truth for comparing the datasets. The DEA results were downloaded from Supplementary Datasets 6 and 13 of Wilson et al. [24]. The DEA results were used as input to TRIPS and PCSF. Genes were considered DEGs at log2 fold change > 0.25 and adjusted p-value < 0.05. For RegEnrich and CaCTS, which require expression data as input, the raw counts were extracted from the Seurat R object downloaded from the cellxgene website according to the author’s instructions https://github.com/p4rkerw/Wilson_Muto_NComm_2022?tab=readme-ov-file. Counts were filtered based on cpm > 1 and transformed using voom. Ensembl IDs were mapped to gene symbols using the mygene package [25].

#### Prostate cancer dataset

We evaluated the different TF prioritization methods on paired single-cell RNA-seq and scATAC-seq data in prostate cancer using the Taavitsainen et al. dataset. The study compared two enzalutamide-resistant LNCaP-derived cell lines (RES-A and RES-B) with the parental DMSO-treated LNCaP prostate cancer cell line as the control to characterize the transcriptional programs mediating drug resistance. Chromatin accessibility profiles were obtained using scATAC-seq. TFs with target genes assigned to the DARs were used as the ground truth for comparing the TF prioritization methods. The DEA results were used as input to TRIPS and PCSF. Genes were considered DEGs at log fold change > 0.25 and adjusted p-value < 0.01. For RegEnrich and CaCTS, which require expression data as input, the raw counts from scRNA-seq were extracted from the GEO database under the accession number GSE168669. Counts were filtered based on cpm > 1 and transformed using the voom function in the R package limma [26].

### 2.2 Biological networks

The human protein-protein interaction (PPI) network was obtained from BioGRID (accessed July 2023) [27], which is derived from manual curation of protein interactions. After retaining only scored edges to filter out weak interactions, the final PPI network contained 15,905 nodes and 329,043 edges. We used two template GRNs for evaluation: DoRothEA and ExTRI. DoRothEA [28] compiles TF-target associations from multiple sources, including ChIP-seq data, literature mining, and motif information. The GRN was filtered for high-confidence edges (confidence levels A and B). The final GRN contained 5,251 nodes and 15,051 unique TF-target edges covering 685 TFs. The second GRN used was the ExTRI resource, which is derived from text mining [29]. ExTRI was filtered for high-confidence TF-target interactions (confidence=“high”), with the final ExTRI GRN containing 4,170 nodes and 18,368 edges.

### 2.3 TRIPS workflow

TRIPS performs a two-step workflow that combines disease module identification on a PPI network with the PCST algorithm on a template GRN (Figure 1). The first step aims to augment the list of DEGs to provide network context by including interactors of the DEGs on the protein level. The second step is used to discover transcriptional regulatory modules derived from the augmented gene list and the signals from experiment-specific DEA results.

**Figure 1.**
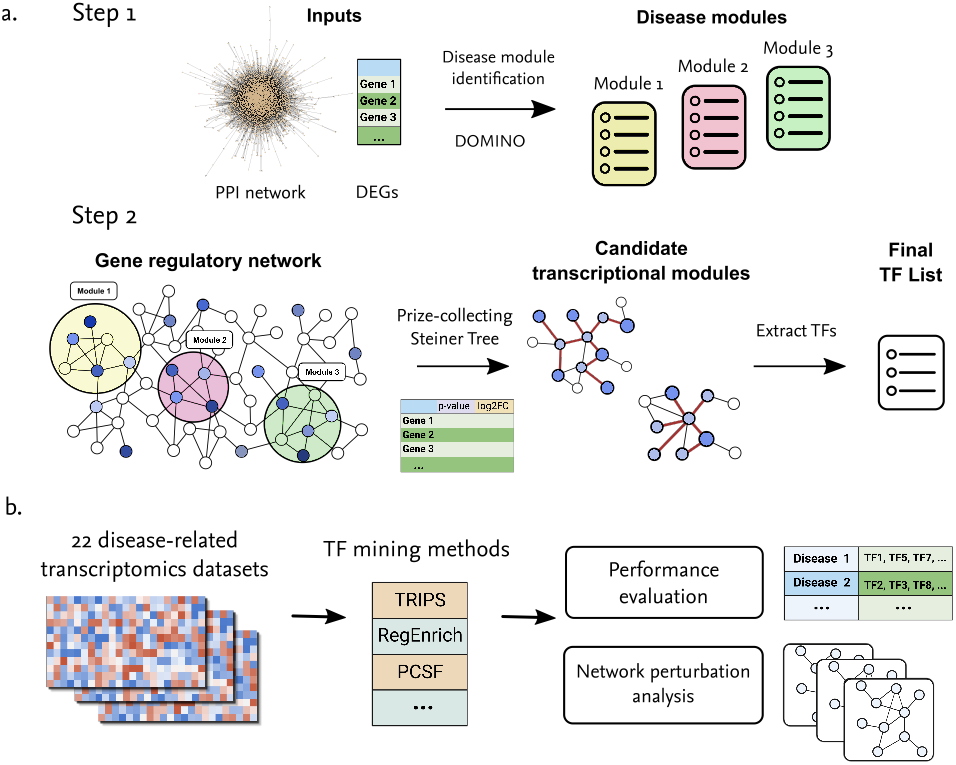
a) Overview of the TRIPS workflow. The inputs are the molecular networks and the DEA results. The first step involves disease module identification using DOMINO, using the list of DEGs as seeds to obtain an augmented list of genes. The second step involves solving the prize-collecting Steiner tree problem to output transcriptional modules. b) Evaluation workflow for the compared TF mining methods.

#### 2.3.1 Disease module identification

Disease modules (“active” regions on a PPI network putatively associated with a disease) are first identified using the Discovery of active Modules in Networks using Omics (DOMINO) algorithm [30]. The rationale for performing this initial step is twofold: 1) TFs exert their function as proteins and not as transcripts, motivating the integration of PPI information. Intricate multilayer control of regulatory programs involves not only direct binding of TFs to a target gene’s regulatory regions, but also the cooperativity between TFs and TF-binding proteins. [31–34]. Furthermore, PPI network analysis has been shown to explain the effects of TF knockout at the gene expression level [35]. 2) Given the generally low abundance of TF expression levels and their possibly short-lived temporal activity, robust detection using gene expression data can be challenging [11, 36, 37]. Thus, augmenting the list of DEGs with other proteins that have been experimentally validated to interact with their protein products can provide further mechanistic context and improve the detection of relevant transcriptional modules.

For each dataset, disease module identification was performed using DOMINO [30]. The DOMINO algorithm was used in the TRIPS workflow because of its superior performance based on a previous benchmarking study of active module identification tools [20]. DOMINO first topologically partitions the PPI network into smaller communities using the Louvain algorithm and subsequently performs enrichment-based tests to identify one or more candidate modules. Module refinement steps that include solving the PCST problem are then performed to obtain the final compact disease modules.

#### 2.3.2 Prize-collecting Steiner trees

Next, we solve the prize-collecting Steiner tree (PCST) problem to identify candidate transcriptional modules. For the task of TF mining, we apply the PCST problem on the GRNs. This step finds coherent high-scoring subnetworks based on the signals derived from the DEA. The PCST problem is a classic *NP*-hard combinatorial problem in graph theory. Given an undirected graph *G* = (*V, E*), where the nodes are assigned non-negative prizes *s* : *V* → ℝ_≥0_ and edge costs *c* : *E* → ℝ_≥0_, the PCST problem finds a subnetwork *G*_0_ = (*V*_0_, *E*_0_) with *V*_0_ ⊆ *V* and *E*_0_ ⊆ *E* having a tree structure that minimizes the following objective function:

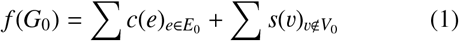

Minimizing *f* (*G*_0_) is equivalent to collecting a set of high-prize nodes while minimizing the cost of the set of edges that are used to connect them. To solve the PCST problem, TRIPS uses dapcstp, which implements a dual ascent, branch-and-bound framework for solving the Steiner arborescence problem and can be used to solve other variants of the Steiner tree problem [38].

The PCST problem is solved for each identified DOMINO module. The node prize assigned to gene v in a disease module is calculated as follows:

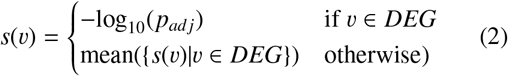

Where *DEG* is the list of DEGs and *p*_*ad j*_ is the adjusted p-value of a gene derived from the experimental results. The DEGs in the identified DOMINO module are assigned prizes using the adjusted p-values derived from the DEA, while other genes in the module are assigned the mean of the prizes of the DEGs that were included in the module. Alternatively, the non-DEGs in the modules could be assigned the minimum score of the DEGs in the module to place higher importance to the DEGs. We further apply a hub penalty to reduce the influence of hubs in the network that decreases the node prizes based on their degree in the backbone GRN:

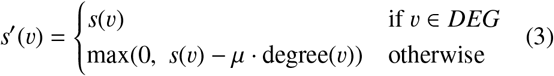

Where µ is the penalty parameter and degree(v) is the node degree of gene v in the GRN *G*. To retain the context specificity from each experiment, nodes that are part of the DEGs were excluded from the hub penalty. The final solution is obtained by merging the identified subnetworks. We set the edge cost to the user-defined *m*th percentile of the adjusted p-values to make the calculation of edge costs adaptive to the score distribution for each dataset. To promote compact, high-scoring subnetworks, we set a high default value of *m* = 50. The hub penalty was set to µ = 0.05.

### 2.4 Compared methods

We compared TRIPS to three baseline methods and four other relevant methods that accept gene expression data with or without a template network as input. All evaluated network-based methods were run using the DoRothEA or ExTRI network as the backbone GRN. The DEGs workflow simply lists all the TFs in the list of DEGs and does not use a backbone network. The DEGs+PCST workflow directly applies the PCST algorithm using the DEA results and skips the disease module identification step. We also ran the popular network inference tool ARACNE [39] that generates a gene regulatory network from expression data by calculating pairwise mutual information and filtering out indirect connections between genes. In the ARACNE workflow, the network was built from the top 5000 edges output by ARACNE on the expression data in the disease condition, followed by extraction of the TF lists.

The Cancer Core Transcription factor Specificity (CaCTS) method infers master transcriptional regulators in a purely data-driven manner [40]. Originally applied to cancer-related datasets, it calculates a score based on the Jensen-Shannon Divergence (JSD), which measures the similarity between two distributions. The final list of master regulators for an experiment is obtained by intersecting the most highly expressed TFs with those having the highest JSD scores. To run CaCTS, we set the parameter denoting the intersection to the top 10%, as the more stringent default value of 5% did not find intersections in the majority of the datasets.

RegEnrich performs differential expression analyses and runs a network inference method to obtain a GRN. Alternatively, a custom network can be provided. Next, it uses enrichment tests to rank and prioritize TFs based on the idea that the targets of TFs in the inferred GRN tend to be differentially expressed. Finally, RegEnrich outputs a list of candidate regulators ranked based on a combined score from the differential expression and enrichment results [41]. To run RegEnrich, we set “DESeq2” as the differential expression analysis method and “GSEA” for enrichment testing, which was found to perform better than the “FET” (Fisher’s exact test) approach. Edge weights in the GRN were set to 1.

The prize-collecting Steiner forest (PCSF) is another approach related to the Steiner tree problem [42]. Given an undirected graph *G* = (*V, E*), where the vertices are labeled with prizes/scores *s* : *V* → ℝ_≥0_ and the edges are assigned costs *c* : *E* → ℝ_≥0_, PCSF aims to find a subnetwork *G*_0_ = (*V*_0_, *E*_0_) with *V*_0_ ⊆ *V* and *E*_0_ ⊆ *E* having a forest structure that minimizes: 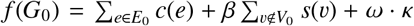 where κ represents the number of trees (components) in the forest solution and can be tuned using the ω parameter. The β parameter adjusts the importance of the node prizes relative to the edge costs. Additionally, a hub penalty to a node v can be applied: *s*^′^(*v*) = *s*(*v*) − *µ* · degree(*v*) with tuning parameter µ. PCSF was originally developed to recover active modules on a PPI network. Similar to TRIPS, the formulation of PCSF directly allows for finding multiple molecular subnetworks. However, PCSF can only handle a single input template network; thus, in contrast to TRIPS, an additional disease module identification step cannot be performed for the task of TF mining. For the b parameter, which represents the prize multiplier for the nodes, we used values *b* = 1, 2. The number of identified subnetworks for the PCSF method should be provided by the user. Thus, for the *w* parameter, which represents the number of trees comprising the forest, we used a range of values *w* = 2, 3, 4, 5, 6, 7. The hub penalty parameter was set to *µ* = 0.05.

The MOGAMUN algorithm is an active module identification method that can use multiplex biological networks [43]. MOGAMUN implements a genetic algorithm that adapts the Non-dominated Sorting Genetic Algorithm II algorithm to networks and optimizes two objectives, namely, the node scores and the density of interactions in the output subnetwork. MOGAMUN can utilize both a template PPI network and a GRN; however, unlike TRIPS and PCSF, the algorithm aims to identify a single connected subnetwork. For each dataset, we run MOGAMUN ten times; in each run, we set `Generations=100` and the default values of `PopSize=100, TournamentSize=2, MinSize=15, MaxSize=50, JaccardSimilarityThreshold=30, CrossoverRate=0.8, MutationRate=0.1, MaxNumberOfAttempts=3, Measure=FDR`, and `ThresholdDEG=0.05`. Results across runs were combined using the `mogamun_postprocess` function [43].

### 2.5 Network generators

Network-based methods should be able to effectively utilize the knowledge encoded in a template network to generate its predictions. Thus, to evaluate the reliability of the compared methods, the TF mining methods were run on perturbed GRNs. For the undirected networks, the original GRNs were perturbed using network generators from the active module identification methods (AMIM) test suite [20], namely, the rewired, expected degree, scale-free, and shuffled networks. Rewired networks are generated by degree-preserving randomization and thus preserve the node degrees. Expected degree networks preserve the node degrees in expectation. The generated scale-free networks have similar overall structure to the original networks but do not preserve node degrees. Shuffled networks are generated by shuffling the gene names.

### 2.6 Gold standards and evaluation metrics

Known disease-associated genes were downloaded from the DISEASES database, an integrated resource primarily derived from text mining [44]. The GEO datasets were mapped to their corresponding DOIDs for evaluation using the DISEASES database based on the dataset descriptions (Supplementary Table S1). The list of human TFs was downloaded from http://humantfs.ccbr.utoronto.ca/download.php (v 1.01) [11]. All human TFs that were associated with a disease were used as gold standard TF lists. To filter for high-confidence TF-disease associations, only entries with a minimum score of 1 in the DISEASES database were considered.

For all analyses, the performance of the evaluated methods were assessed based on precision, number of recovered disease-related TFs, and the solution size. For the related methods TRIPS and PCSF, solution size could also serve as an indicator of the quality of the identified subnetwork because larger subnetworks suggest that nodes with high prizes are located in a coherent region in the network. To evaluate the similarities of the solutions between different methods, we calculated the Jaccard similarity between two TF sets *A* and *B*: Jaccard 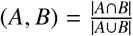. The one-sided Mann-Whitney-U test in the scipy package was applied to determine significant differences in performance when using the perturbed GRNs [45]. Differences were considered statistically significant at *p* < 0.05.

## 3 Results

### 3.1 Evaluation results

Comparison with baseline methods. For our evaluation on the 22 disease-specific datasets, we first compared TRIPS to baseline methods. As shown in Figure 2, TRIPS was found to perform better than the DEGs workflow as well as the ARACNE workflow, which infers the network from gene expression data alone. Furthermore, applying the PCST algorithm directly to the list of DEGs (DEGs+PCST) increases the precision of identifying disease-relevant genes and TRIPS performed even better. The improved performance of TRIPS compared to these two baseline workflows indicate that the implemented strategy provides more biological context through the PPI network. Furthermore, while the DEGs+PCST workflow also showed high precision, TRIPS showed better results based on network perturbation analysis (Figure 2b and Supplementary Figure S1). In addition, using DOMINO in the TRIPS workflow outperformed the popular disease module identification method DIAMOnD [46], both in terms of precision and reliability based on network perturbation analysis (Supplementary Figure S2). Overall, the results indicate that using DOMINO in the disease module identification step provides better molecular context for deriving candidate TFs.

**Figure 2.**
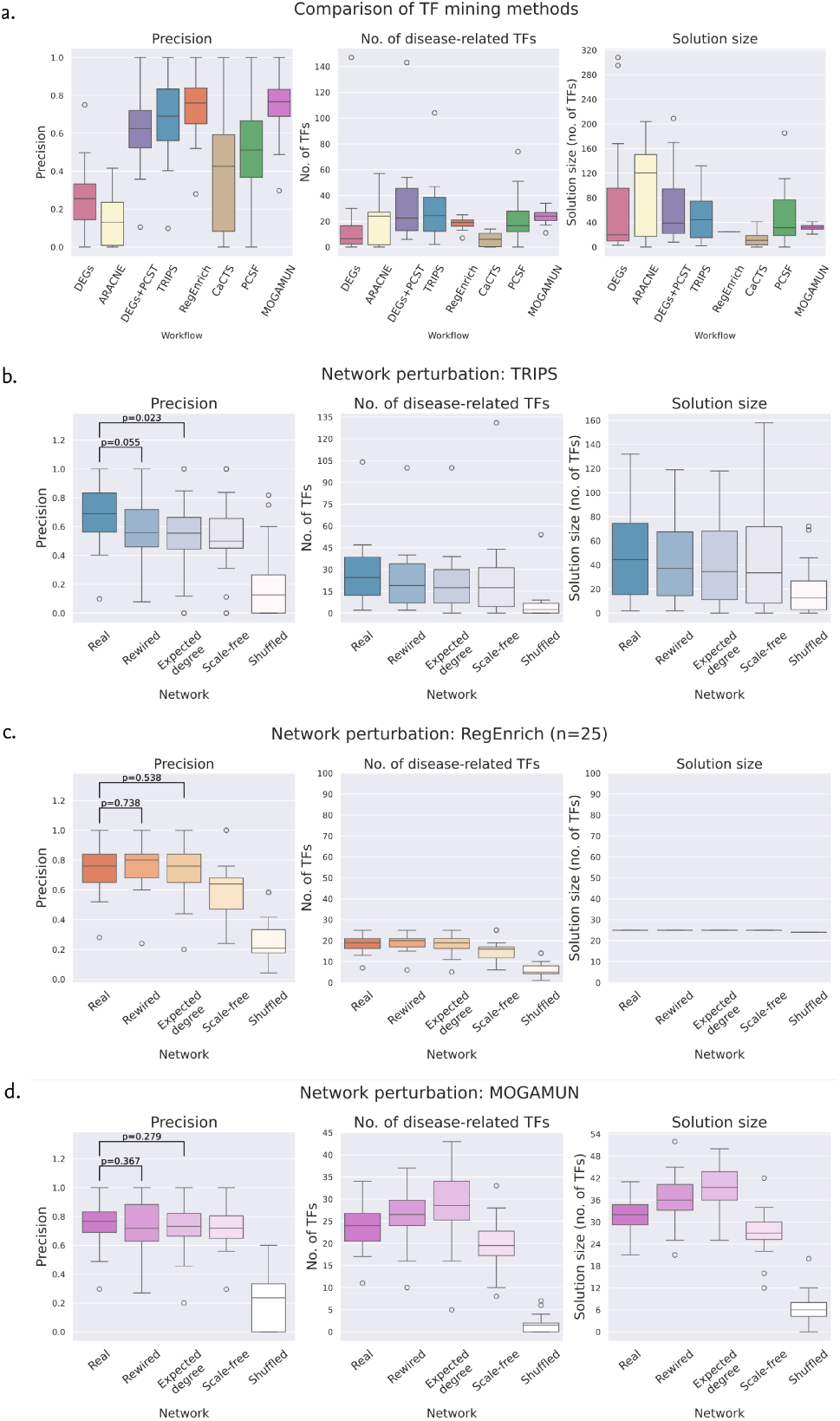
a) Comparison of TF mining methods from baseline methods and network-based methods using DoRothEA as the template GRN. Similar results for PCSF were obtained using other parameters. b) Network perturbation analysis results for TRIPS. c) Network perturbation results for RegEnrich considering the top 25 TFs. Similar results were obtained at increasing cutoff values the top 15 and 50 TFs (Supplementary Figure S3). While RegEnrich showed higher precision in identifying disease-associated TFs, results of network perturbation analysis showed no drop in the performance when using the rewired and expected degree networks. d) Network perturbation results for MOGAMUN. MOGAMUN also showed higher precision than TRIPS; however, it identified higher numbers of diseases-associated TFs when using the rewired and expected degree networks. On the other hand, TRIPS showed good precision with a decrease in performance metrics upon using rewired and random networks.

#### 3.1.1 Comparison with other TF mining methods

Previous findings have highlighted the importance of performing network perturbations to demonstrate the reliability of tools that utilize a reference network, typically a PPI network [20]. Most existing methods were found to primarily generate predictions based on node degrees, indicating that the tools did not rely on the biological knowledge encoded in the network. The above results indicate that the network structure of the template GRN is important for the performance of TRIPS in identifying candidate TFs. As shown in Figure 2a, TRIPS showed higher precision compared to PCSF and the data-driven CaCTS method, but RegEnrich and MOGAMUN showed the highest precision among the compared methods. However, based on network perturbation analysis, RegEnrich returned no difference in precision when using rewired (p=0.816) and expected degree (p=0.656) networks, which have similar structure to the original GRN, even though precision considerably decreased when using the scale-free and shuffled networks (Figures 2b - 2c). For all methods, however, performance significantly decreased when using shuffled networks. For MOGAMUN, we similarly observe no significant difference in precision for both the rewired and expected degree networks (Figure 2d), and even observe an increase in both the number of recovered TFs and solution size. The precision of MOGAMUN in identifying disease-related TFs evidently dropped only when using the shuffled network. On the other hand, only TRIPS showed a decrease in precision when using the rewired network as the template network (p=0.055), and a statistically significant difference when compared against the expected degree network (p=0.023). None of the other evaluated methods showed a significant drop in precision, even when using the expected degree network. While precision was low for PCSF, performance slightly decreased when using perturbed networks (Supplementary Figure S1). TRIPS solutions obtained from the real template networks had low overlap with those obtained with the perturbed networks (Supplementary Figure S4). In general, TF sets output by TRIPS had low overlap with those of the other methods (Supplementary Figure S5).

#### 3.1.2 Effect of the backbone network

We additionally evaluated the network-based TF prioritization workflows using a different template GRN, ExTRI, which is derived from text mining. For TRIPS, RegEnrich, and PCSF, their overall relative performance was consistent with those of the DoRothEA results (Supplementary Figure S6). However, using DoRothEA as the template GRN showed higher prediction performance and better network perturbation results, especially when compared to the rewired network, highlighting the greater reliability of GRNs primarily obtained from experimental evidence. Interestingly, for MOGAMUN, using the ExTRI GRN as backbone led to higher precision compared to DoRothEA. However, this could indicate literature bias, since network perturbation results using ExTRI remained poor. In particular, we observe no significant difference in precision between real, rewired, and expected degree networks (Supplementary Figure S7). In addition, similar to the network perturbation results obtained using DoRothEA, the number of disease-associated TFs that were recovered by MOGAMUN were higher for the rewired and expected degree networks, even though performance dropped in scale-free and shuffled networks.

#### 3.1.3 Integrating PPI and GRN information

Note that compared to the other methods, TRIPS requires both PPI and GRN information as input. Unlike MOGAMUN and TRIPS, RegEnrich and PCSF do not provide a natural framework for analyzing both types of data, as only a single input network is considered. An alternative is to combine the PPI and GRN into an integrated network. However, using the PPI+GRN network as input to the DEGs+PCST, RegEnrich, and PCSF workflows led to a decrease in performance for all three methods (Figure 3). Simply concatenating the edges from PPI and GRN introduces more edges, which could decrease the overall network diameter as well as introduce false positive interactions, leading to poor performance.

**Figure 3.**
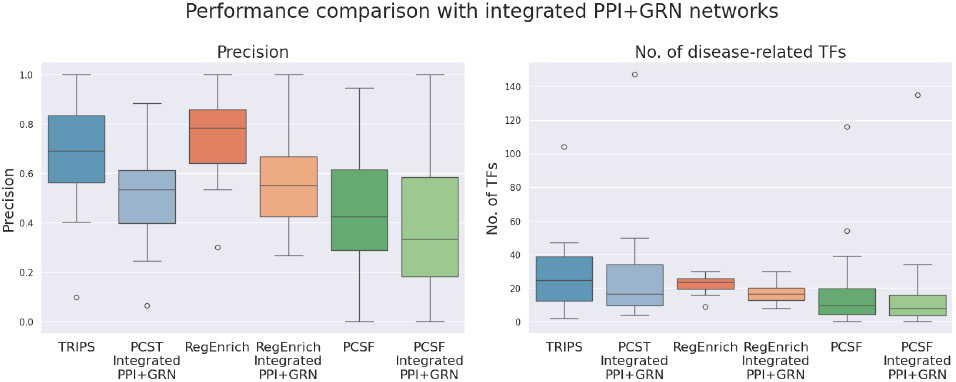
TF mining workflows using integrated PPI+GRN networks. Notably, the two-step TRIPS workflow performs better than applying PCST on an integrated PPI+GRN network or the GRN backbone alone.

### 3.2 Application to Alzheimer’s disease

Alzheimer’s Disease (AD) is a neurodegenerative disorder that is associated with memory loss and progressive decline of cognitive function. AD is common in elderly people and affects around 50 million people worldwide [47]. Key molecular features of AD include extracellular accumulation of amyloid plaques, predominantly the amyloid β (Aβ) peptide, as well as the deposition of neurofibrillary tangles. Multiple pathogenic processes have been linked to AD, including the major contribution of microglia, resident brain myeloid cells that play critical roles in neuroimmunity and homeostasis [48, 49]. Microglia may exert damaging effects in the brain by inducing the expression of pro-inflammatory cytokines.

To demonstrate the practical utility of TRIPS, we applied it to two AD datasets: snRNA-seq putaminal microglia from Xu et al. [22] and paired snRNA-seq and snATAC-seq data cortical microglia from Morabito et al. [23]. In the [22] dataset, homeostatic microglia (Micr-0) refer to microglia that have not yet been activated, while activated microglia (Micr-1) are microglia that have switched to a proinflammatory state, leading to the release of cytokines and activation of other immune cells [50]. The two microglial clusters Micr-0 and Micr-1 were distinguished by elevated *P2RY12* expression in Micr-0 and reduced *P2RY12* expression in Micr-1; upregulation of the inflammation- and antigen presentation-related *AIF1, CD14*, and MHC II genes in Micr-1; and pathway analysis results indicating elevated enrichment of immune response genes in Micr-1. The TF sets output by each tool are shown in Supplementary Table S2. While the tools returned generally different results, TRIPS, RegEnrich and MOGAMUN showed the most similar TF sets (Supplementary Figure S8).

In homeostatic microglia in the AD putamen, enriched pathways include inflammation-related pathways (as in cytokine production regulation and cytokine-mediated signaling in subnetworks 2 and 3, and regulation of inflammatory response in subnetwork 1), regulation of protein binding (as in subnetwork 3), and apoptosis-related pathways (as in subnetwork 3) (Figure 4). This is consistent with the findings of Xu et al. [22] for homeostatic microglia in AD and may reflect the engagement of programs that are adaptive to the chronic inflammatory state in AD [51]. In activated microglia, death-related pathways are enriched in the subnetworks 1 and 2, including regulation of neuron apoptotic process in subnetwork 1, while signal transduction by p53 is enriched in subnetwork 3 (Figure 5). Furthermore, inflammatory pathways, such as production and response to cytokines, are enriched in subnetworks 1 and 2. Moreover, oxidative stress response is enriched in subnetwork 2. These findings are again consistent with those of [22, 23] and the described actions of activated microglia, including response to and release of inflammatory mediators [52], and amyloid beta binding and clearance [53–55]. Moreover, neuronal death regulation may constitute the neuron-microglia crosstalk [56] such as during the phagocytosis of apoptotic neurons in neurodegeneration, seen in the activated––not the homeo-static––microglial phenotype [57].

**Figure 4.**
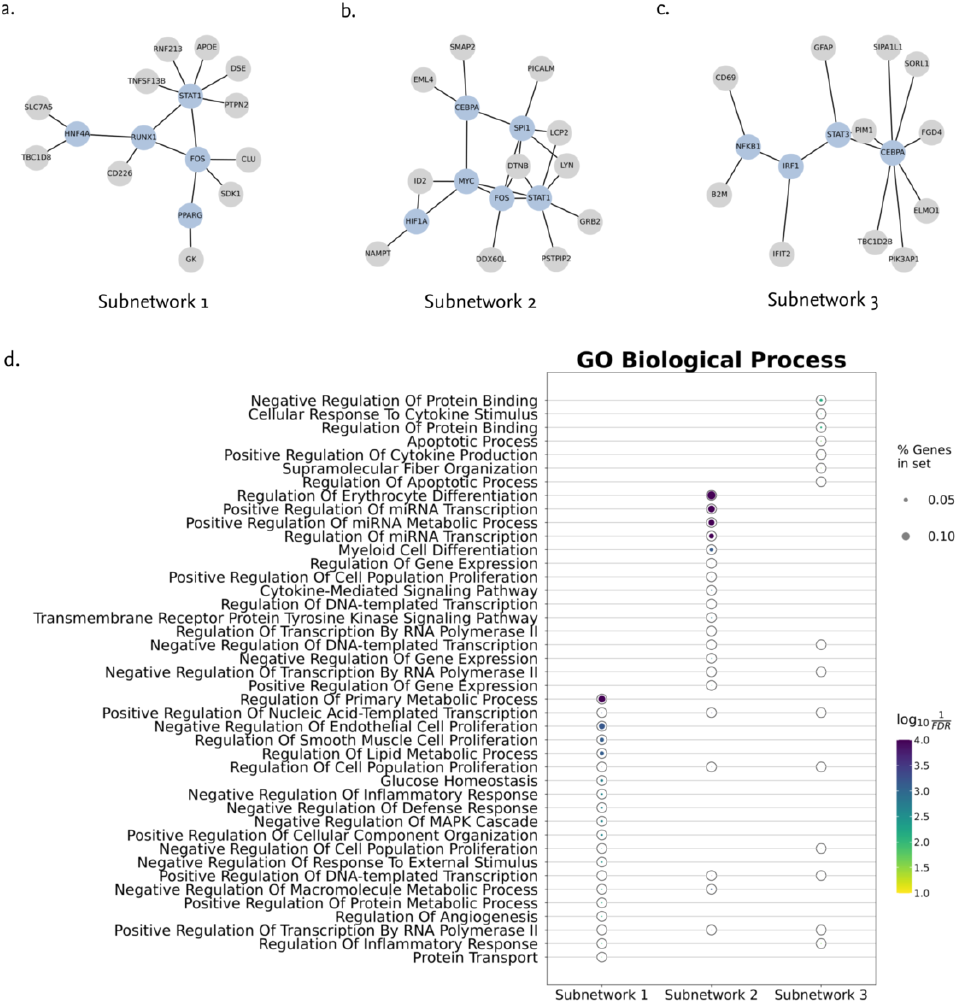
a-c) Top three largest induced subnetworks from homeostatic microglia (Micr-0). TFs are colored in blue, while target genes are colored in gray. d) Enriched GO Biological Process terms. Subnetwork 1 was found to be enriched in terms related to interferon signaling. Subnetwork 2 was enriched in path-ways related to catabolism of amyloid precursor protein. Subnet-work 3 was enriched in terms related to inflammatory response, including regulation of cytokine and response to interleukins.

**Figure 5.**
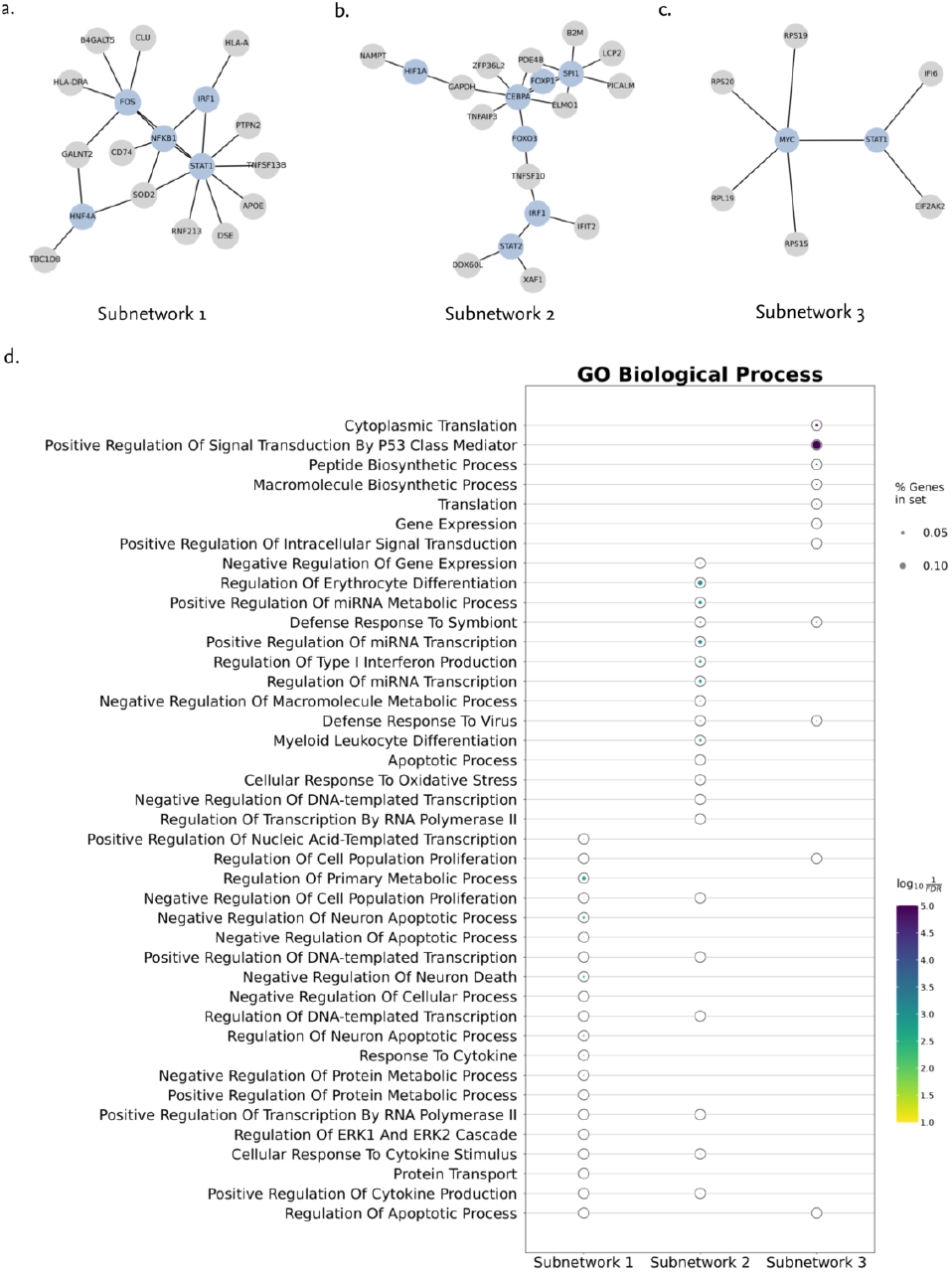
a-c) Induced subgraphs from the three largest TRIPS modules from activated microglia in the Xu et al dataset. TFs are colored in blue, while target genes are colored in gray.d) enriched pathways. Subnetwork 1 was found to be enriched in terms related to neuronal death. Subnetwork 2 was enriched in pathways related to miRNA regulation and cytokine signaling. Subnetwork 3 was also enriched in terms related to P53 signaling and cell proliferation.

In the Morabito et al. dataset, only snRNA-seq data were used as inputs and the resulting TF lists were vali-dated using independently profiled snATAC-seq data from the same samples. In this dataset, the use of both snRNA-seq and snATAC-seq makes the determination of clusters that are homeostatic and activated microglia difficult (as they have different clusters in the two modalities) − so all microglia are lumped together into one major cluster in the analysis. Relying on DEGs alone identifies only two TFs that have significantly differentially enriched TF motifs between AD patients and healthy controls based on snATAC-seq profiling. On the other hand, out of the 12 candidate TFs identified by TRIPS, seven were supported by the snATAC-seq results, namely *NFKB1, FOS, STAT1, CEBPA, GATA3, TRPS1*, and *TFAP2C*, corresponding to a precision of 0.583. We compared this to results obtained using the other methods. RegEnrich identified 15 out of 25 TFs that were supported by the chromatin accessibility results, corresponding to a comparable precision of 0.6. These include *RUNX1, MEF2C, ESR1, STAT1, CEBPA, HNF4A, TFAP2C, GATA3, FOS, CTCF, TAL1, MITF, SPI1, FOXA1, GATA2, SOX2, TP53, FOXP1, MEF2A*, and *ETS1*. MOGAMUN identified 23 out of 30 TFs (*TFAP2C, GATA2, SPI1, MYC, SMAD3, CTCF, GATA3, FOXA1, POU2F1, NFATC2, STAT3, TP53, YY1, RARA, NFKB1, ESR1, VDR, FOS, CEBPA, ETS1, JUN, FOXP1*, and *STAT1*), with the highest precision of 0.77. CaCTS identified 15 out of 68 TFs (*ELF1, NR3C1, POU2F2, HIF1A, RBPJ, ETV6, NFATC2, SMAD3, CREB1, NFKB1, NRF1, FOXK2, REL, STAT3* and *PBX3*) that are consistent with the ATAC-seq results, corresponding to a precision of 0.22. On the other hand, PCSF did not identify a coherent subnetwork.

There are two large transcriptional modules identified by TRIPS in the Morabito et al. [58] dataset, which profiled the cortical AD microglia. Notable pathways enriched include cell death regulation, cytokine response, endoplasmic reticulum (ER)–associated protein degradation and ER stress response in subnetwork 1; and cytokine response in subnetwork 2. These findings are consistent with but expand on the pathway analysis results of the microglial cluster differential gene expression analysis in [23]. While cell death regulation and cytokine response are a recurring theme across microglial datasets in AD here and elsewhere [54, 59–6 our TRIPS analysis supplements this with ER-related protein degradation and stress response. These pathways can result in or get disrupted from the accumulation of misfolded proteins, such as amyloid beta and hyperphosphorylated tau, which could lead to the unfolded protein response [63].

Notable AD-associated TFs identified by TRIPS in both AD datasets include *GATA3, CEBPA, FOS, SPI1, STAT1*, and *NFKB1. GATA3*, which is known to promote inflammation, is activated in phagocytic microglia [58, 64, 65]. *FOS* is involved in microglial development and in promoting the inflammatory “disease-associated microglia” cell state [66, 67]. *NFKB1* is a canonical inflammatory effector molecule that has multiple binding sites in the promoter regions of the genes involved in amyloidogenesis [48, 68]. *STAT1* signaling mediates memory loss by inhibiting the expression of N-methyl-D-aspartate receptors, which play crucial roles in synaptic function [69]. *CEBPA* is a co-factor of *SPI1*, an established regulator of microglial development that influences antigen presentation and phagocytosis functions [70– In AD models, APOE phospholipids lead to STAT1 activation, which in turn promotes the inflammatory state of microglia [75]. Differential activity of these TFs in AD could thus mechanistically underlie disease pathophysiology, consistent with the inflammatory, amyloidogenic, and synapto-pathic features of AD [76**–**78].

### 3.3 Evaluation on diabetic kidney disease and prostate cancer datasets

We then evaluated the methods using two additional datasets with paired sc/snRNA-seq and sc/snATAC-seq data. The DKD dataset comprised 15 different cell types in the nephron of the diabetic kidney; disease expression was compared to controls. TRIPS showed high precision with respect to the list of TFs showing differential motif enrichment derived from chromatin accessibility data (Figure 7a). The network-based methods TRIPS, RegEnrich, MOGAMUN, and PCSF performed better than the data driven method CaCTS, also demonstrating the advantage of using a backbone GRN in analyzing single-cell gene expression data. We additionally evaluated the different methods on a prostate cancer dataset that compares the transcriptional programs of two drug-resistant LNCaP cell lines (RES-A and RES-B) to the parental prostate cancer LNCaP cell line. While the precisions in identifying candidate TFs supported by the paired sc/snATAC-seq data was low, TRIPS outperformed the other methods for this dataset in both cell lines (Figure 7b).

**Figure 6.**
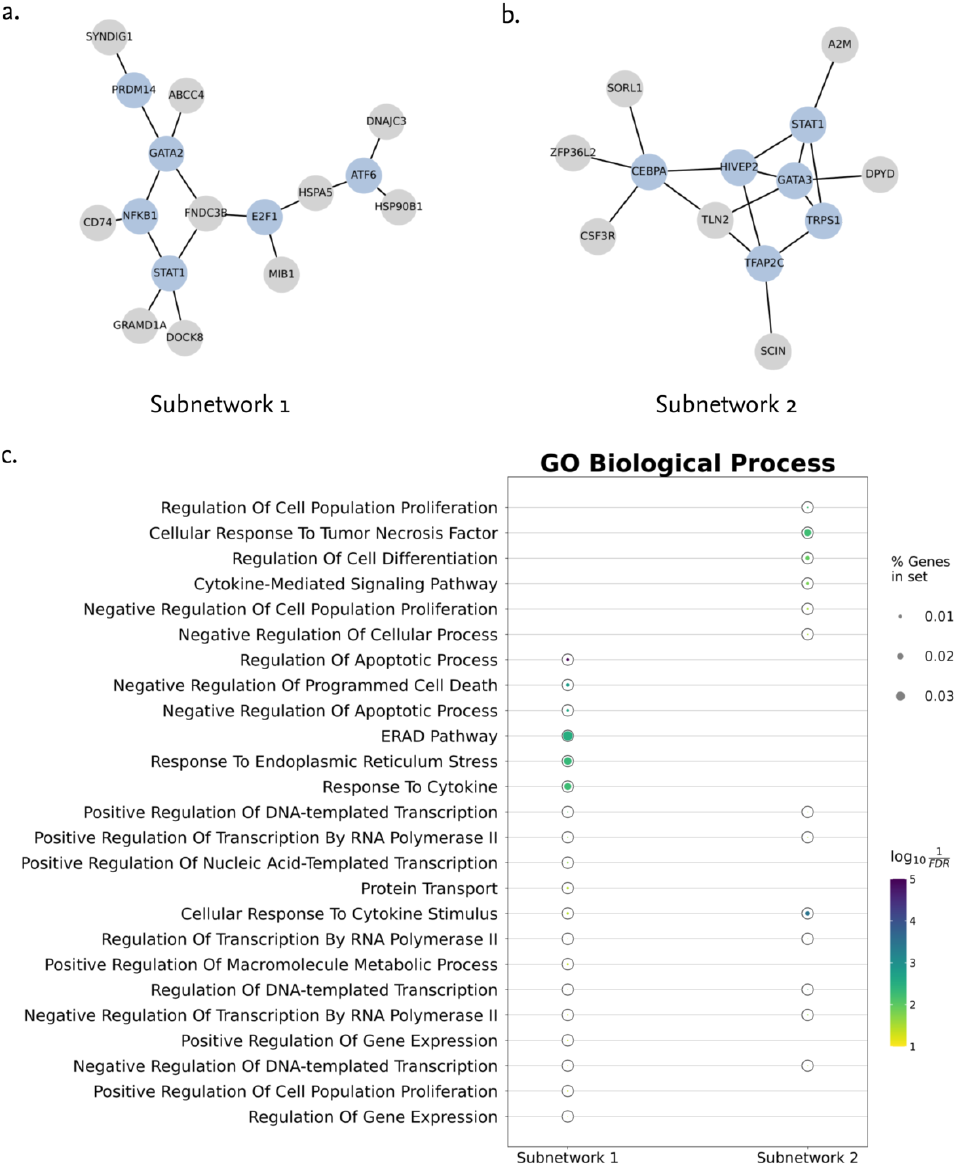
a-b) Induced subgraphs of the two largest solutions obtained from the microglia cell type in the Morabito et al. dataset. TFs are colored in blue, while target genes are colored in gray. c) Enriched GO Biological Process terms. Subnetwork 1 was found to be enriched in terms related to ER stress. Subnetwork 2 was enriched in terms related to cytokine signaling and cell proliferation.

**Figure 7.**
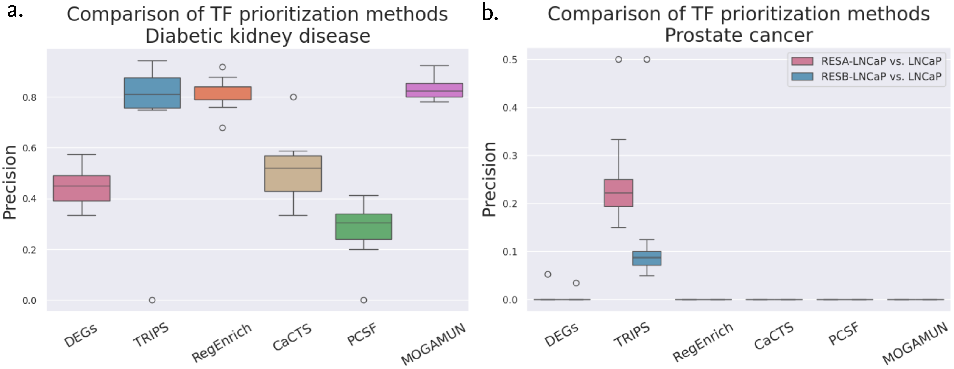
Precision of TF sets with respect to enriched TF motifs in DARs based on chromatin accessibility data in a) the DKD dataset across 15 cell types and b) the prostate cancer dataset of [79] across 13 cell clusters determined by scRNA-seq for the RES-A vs. control and RES-B vs. control comparisons.

## 4 Discussion

TRIPS combines molecular network information from the protein and transcriptional levels in a biologically principled manner to infer candidate TF regulators. Biologically, the final subnetworks obtained by the PCST step in the TRIPS workflow correspond to local neighborhoods of the GRN that have the strongest perturbation signals with respect to the phenotype being analyzed and thus are suitable for prioritization tasks. Using prior knowledge also facilitates interpretability of the results. We demonstrate that TRIPS identifies known disease-associated regulators with high precision using a range of disease-associated transcriptomics datasets. Evaluation on single-cell gene expression datasets also demonstrates that backbone GRNs can be used to provide network context on DEGs that are consistent with known disease pathomechanisms. Moreover, network perturbation analysis results show that TRIPS can more effectively utilize the structure of the biological networks in comparison with other network-based approaches.

The importance of network perturbation analysis has been discussed in a recent study on active module identification tools, which showed that most existing methods generate solutions primarily based on node degrees [20]. In particular, DOMINO was found to outperform other active module identification methods and output more biologically meaningful solutions, consistent with our own evaluation across the 22 datasets in comparison with DI-AMOnD. The algorithmic principle behind DOMINO is to first topologically partition the PPI network into multiple refined modules. Mechanistically, this allows for separate regions of the interactome to be perturbed simultaneously. On the other hand, the majority of other methods assume and effectively detect a single module representing the candidate pathological mechanism. The idea that dysregulation in complex disease conditions can be attributed to multiple molecular mechanisms occurring at the level of molecular networks has been previously highlighted [80, 81]. The flexibility of the PCST algorithm, which relies on assigning prizes to genes by combining one or more layers of evidence, further allows for fine-tuned biological signals to find context-specific subnetworks in addition to using the network structure.

Microglial phenotypes contribute to AD disease states and have been suggested as candidates for precision therapeutics. We demonstrate the use of TRIPS using microglial AD datasets, where the method identified transcriptional regulators that are involved in neurodegeneration and inflammation, consistent with current knowledge of AD pathophysiology. TRIPS also provides a flexible framework to integrate information derived from multiple types of evidence by combining the gene scores derived from omics data or other sources. With the continuous increase in the quality and quantity of data generated from omics technologies, we expect the increased reliability of reference networks and consequently increased applicability of network-based methods such as TRIPS. This type of analysis can be particularly desirable in cases where novel conditions are examined and the sample sizes may be limited. Furthermore, since TRIPS finds transcriptional modules as an intermediate step, it facilitates interpretability. Compared to RegEnrich and CaCTS, methods such as TRIPS and PCSF directly aim to find coherent and mechanistic subnetworks. We further evaluated the methods in two disease contexts, namely, the diabetic kidney and drug resistance in prostate cancer, across multiple cell types or cell clusters using paired scRNA-seq and scATAC-seq data. In general, TRIPS showed greater concordance with the chromatin accessibility results compared to the other methods, indicating the effectiveness of the approach.

Nevertheless, the current study has several limitations. The outputs of network-based methods are influenced by the quality of the backbone networks [82], emphasizing that careful selection of backbone networks or filtering for high-confidence edges is important when performing network-based analysis. However, as more molecular data are generated, including tissue- and cell type-specific data, we expect the increasing reliability of template molecular networks and higher confidence annotation scores of TF-target associations across different biological contexts and the greater applicability for network-based tools. Future work for in-depth characterization of disease mechanisms in complex human diseases involves datasets where multiple tissues or cell types are systematically profiled from the same set of patients.

## 5 Conclusions

TRIPS utilizes template GRNs and PPI networks to contextualize experiment-specific gene expression signals and prioritize disease-associated candidate TFs. Using a range of disease-specific datasets, we demonstrated the high precision of TRIPS in identifying relevant regulators and its reliability through network-based perturbation analyses. Overall, our systematic evaluation of the TF prioritization methods demonstrate the utility of TRIPS in analyzing complex human diseases.

## 6 Availability

All disease-specific gene expression datasets used in the evaluations are publicly available and are listed in Supplementary Table S1. TRIPS is available on GitHub: https://github.com/scibiome/trips.

## 8 Author contributions statement

GG: Conceptualization, Formal analysis, Methodology, Software, Writing - original draft. BAL: Validation, Methodology, Writing - original draft. DBB: Conceptualization, Methodology, Supervision, Writing - Review & Editing. TK: Conceptualization, Methodology, Supervision, Writing - Review & Editing.

