## Supplementary Material for "Identification of transcriptional regulators using a combined disease module identification and prize-collecting Steiner tree approach"

### I. SUPPLEMENTARY FIGURES

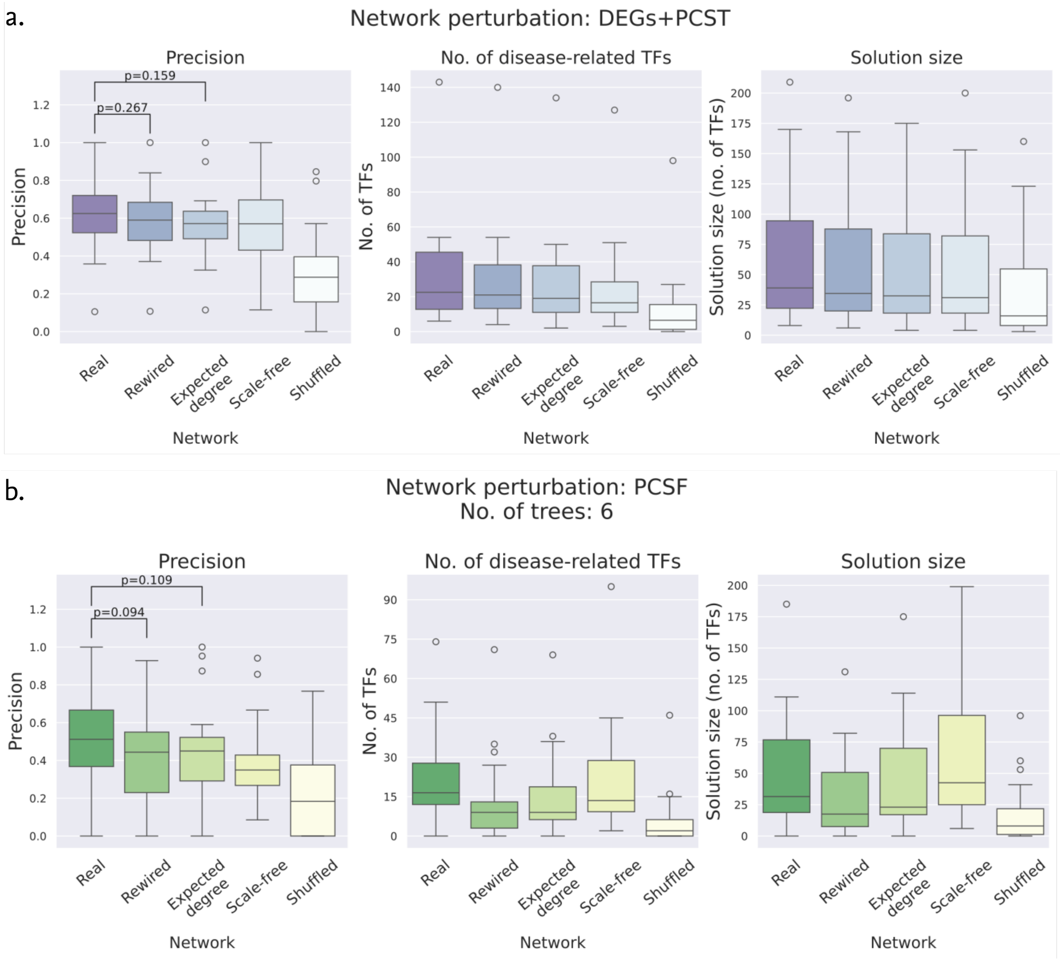


FIG. S1: Network perturbation analysis results for the a) DEGs+PCST and b) PCSF workflows. The precision of the decrease when using rewired or random GRNs, whereas PCSF showed a drop in performance.

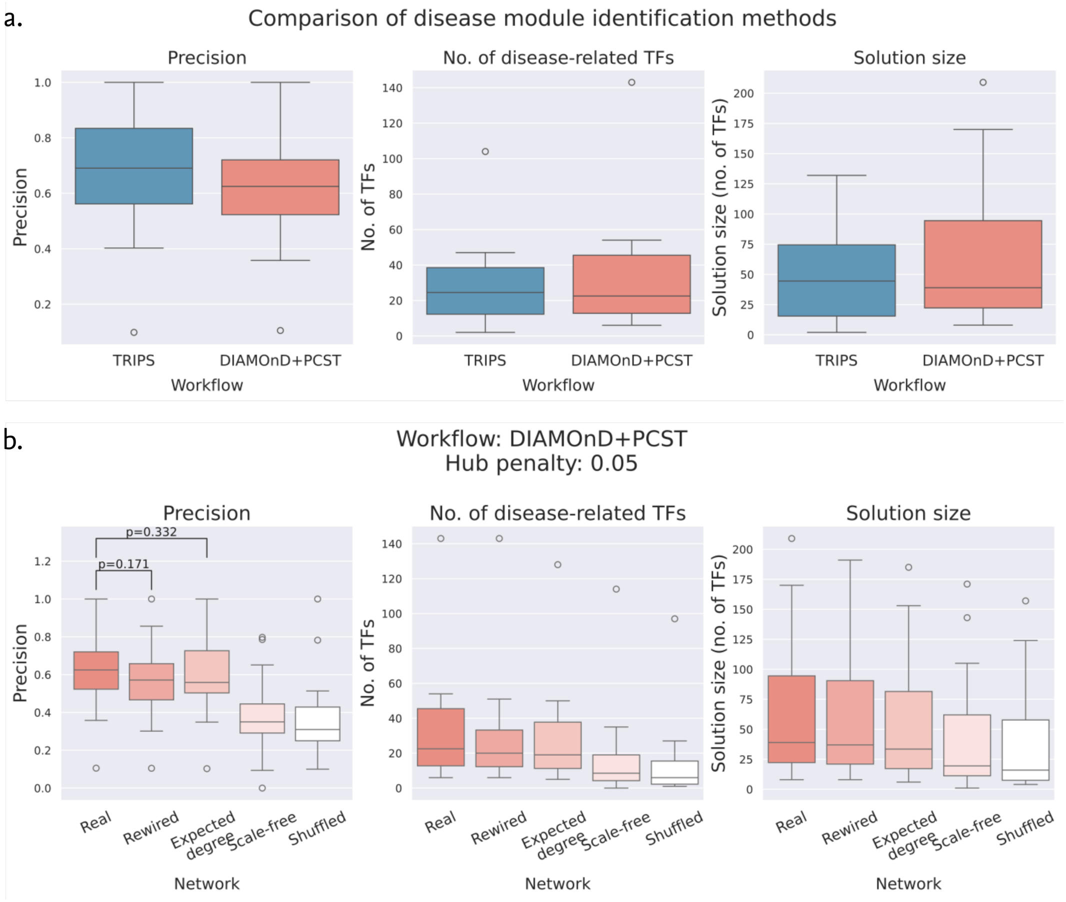


FIG. S2: Comparison of disease module identification methods used in the TF mining workflow. Using DOMINO in the disease module identification step (TRIPS workflow) showed better performance than the DIAMOnD+PCST workflow in terms of precision and network perturbation results. Starting from a list of seed genes, DIAMOnD performs random walks along the PPI network to retrieve genes that exhibit strong connectivity to the seeds in the growing module across iterations of the algorithm [S1].

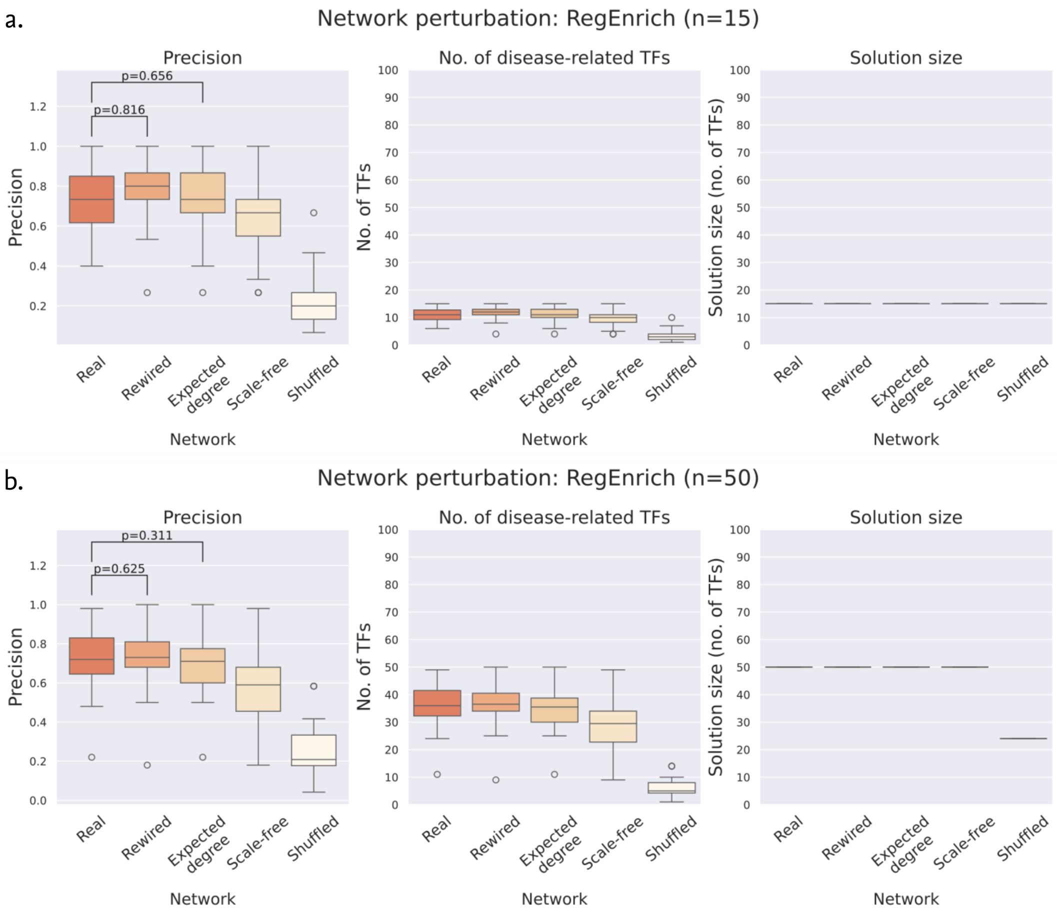


FIG. S3: Network perturbation results for RegEnrich for the a) top 15 regulators and b) top 50 regulators.

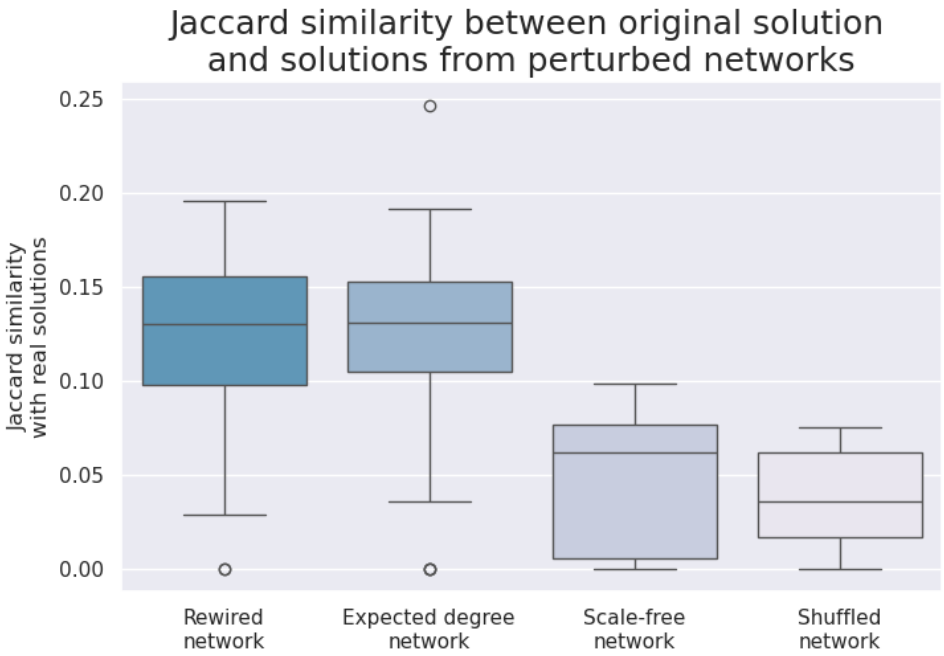


FIG. S4: Jaccard similarities between TRIPS solutions from the original networks and perturbed networks across the 22 datasets.

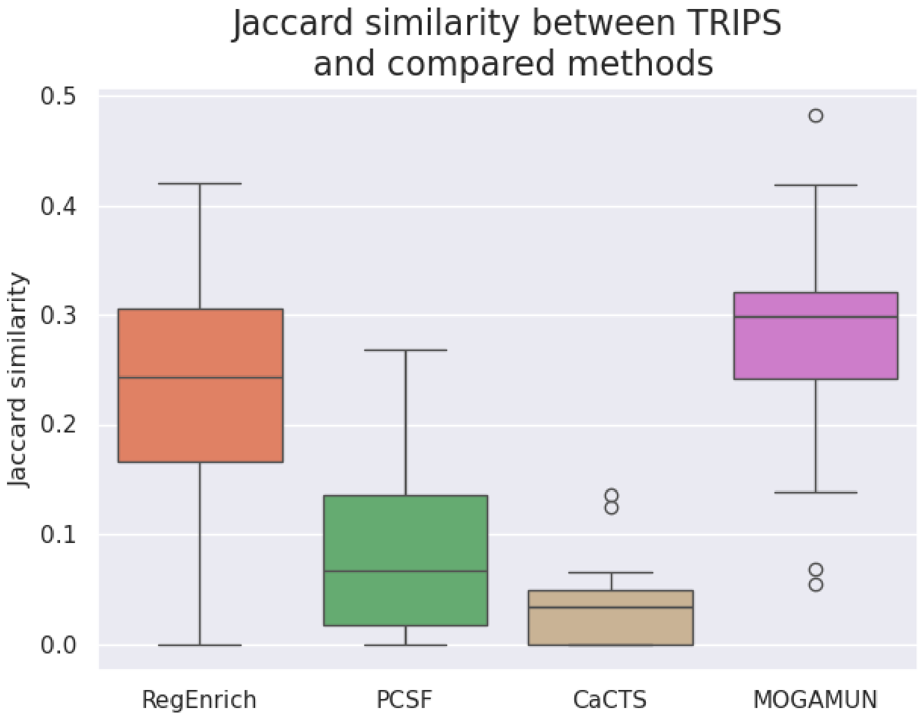


FIG. S5: Jaccard similarities between TRIPS solutions and solutions from other methods across the 22 datasets.

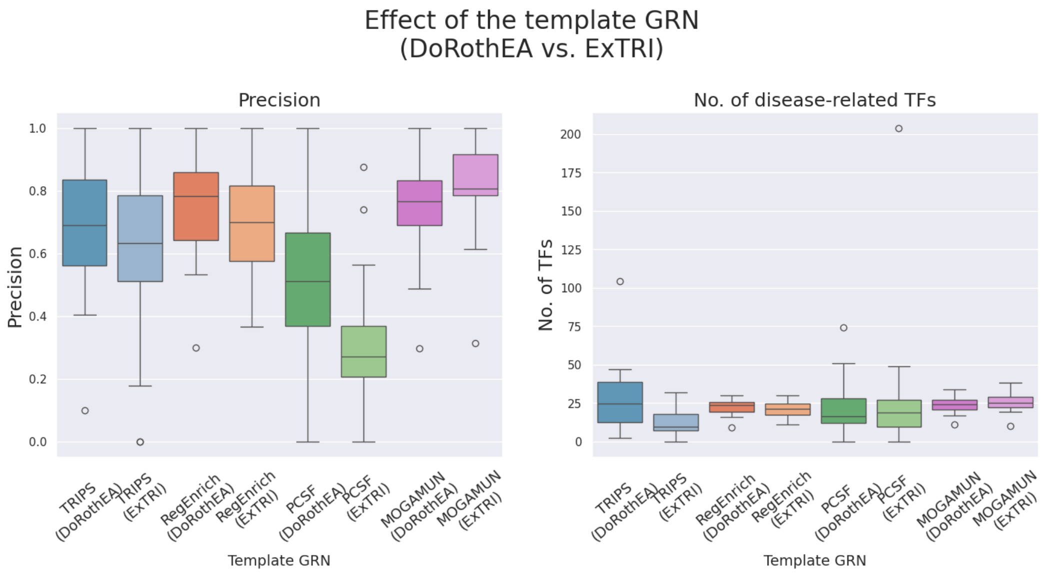


FIG. S6: Evaluation of TF mining methods using ExTRI as the backbone GRN.

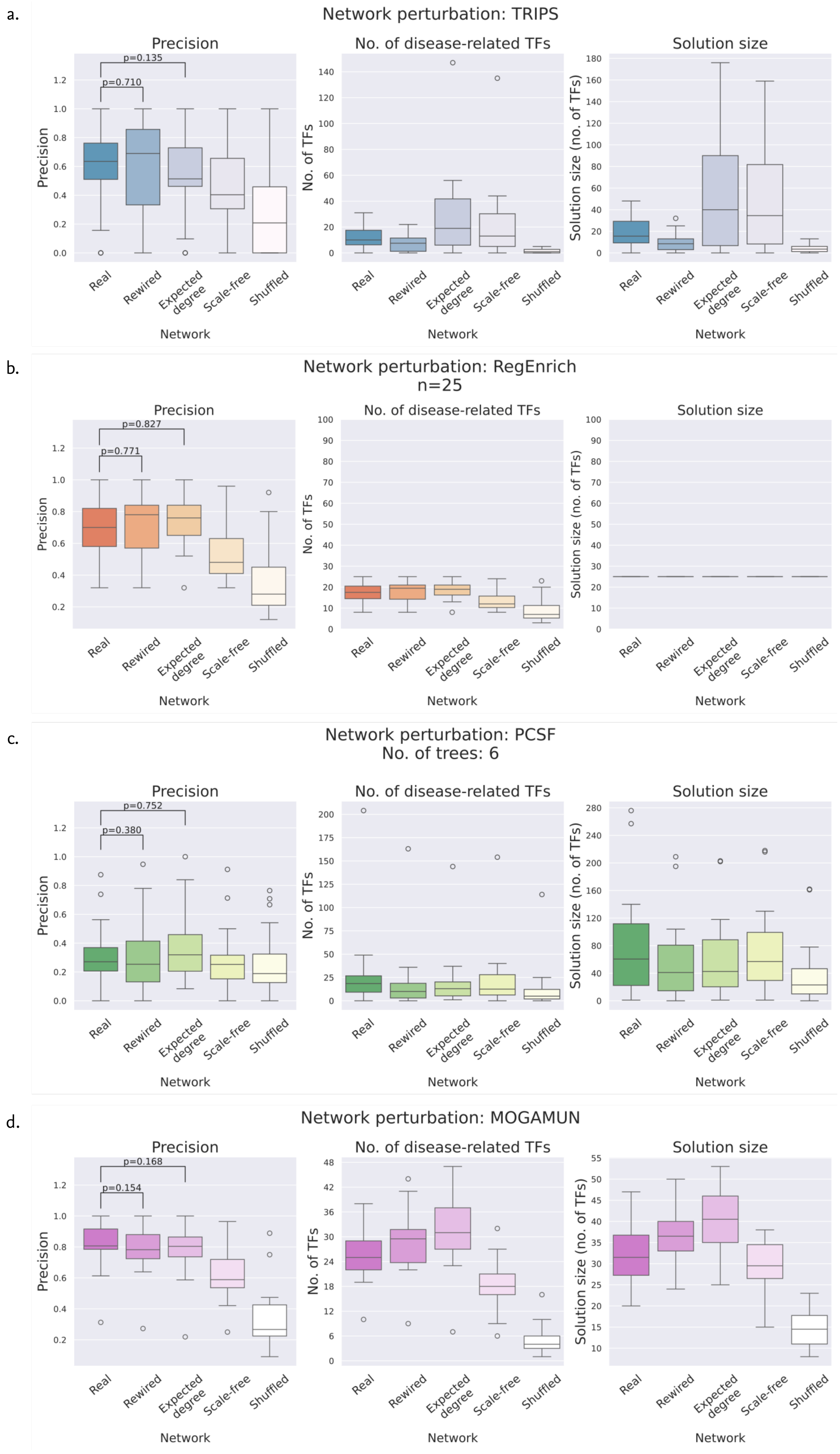


FIG. S7: Results of network perturbation analysis for TRIPS, RegEnrich, PCSF, and MOGAMUN using the ExTRI network.

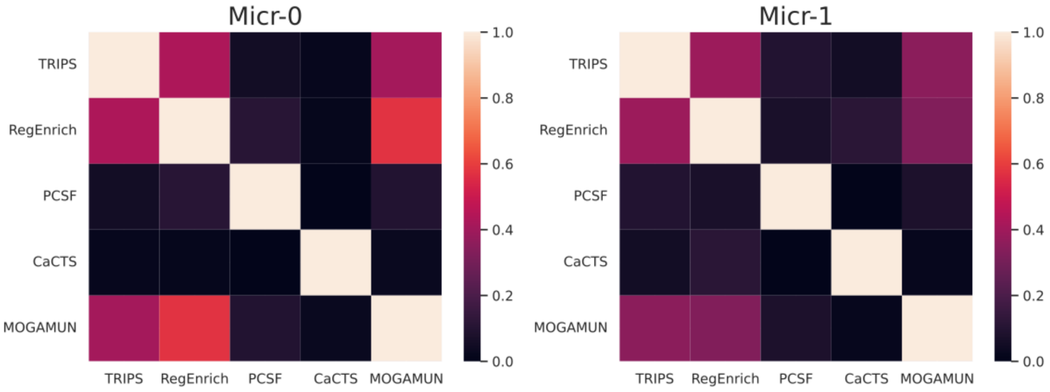


FIG. S8: Heatmap of pairwise Jaccard similarities of the TF sets output by the compared TF mining methods.

### II. SUPPLEMENTARY TABLES

TABLE S1: Disease-associated datasets used for evaluation of TF mining methods.

| GEO dataset | Disease | Tissue |
|---|---|---|
| GSE159984 | Type 2 diabetes | pancreas |
| GSE122459 | Systemic lupus erythematosus | PBMCs |
| GSE72509 | Systemic lupus erythematosus | whole blood |
| GSE97263 | Systemic lupus erythematosus | peripheral blood |
| GSE135251 | Non-alcoholic fatty liver disease | liver |
| GSE126848 | Non-alcoholic fatty liver disease | liver |
| GSE83687 | Inflammatory bowel disease | colon |
| GSE93624 | Inflammatory bowel disease | small intestine |
| GSE137344 | Inflammatory bowel disease | small intestine |
| GSE121212 | Psoriasis | skin |
| GSE95587 | Alzheimer's disease | fusiform gyrus (cortex) |
| GSE125050 | Alzheimer's disease | brain |
| GSE172381 | Preeclampsia | endometrium |
| GSE75011 | Asthma | Blood PBMCs |
| GSE159225 | Multiple sclerosis | leukocytes |
| GSE47462 | Breast cancer | breast mammary tissue |
| GSE57148 | Chronic obstructive pulmonary disease | lung |
| GSE136371 | Cystic fibrosis | blood |
| GSE143323 | Dermatomyositis | skeletal muscle |
| GSE77314 | Hepatocellular carcinoma | liver |
| GSE138614 | Multiple sclerosis | brain |
| GSE86356 | Muscular dystrophy | Bicep muscle |

TABLE S2: TF sets output by the TF prioritization tools for homeostatic and activated microglia in the Xu et al. dataset.

| **Method** | **Micr-0** | **Micr-1** |
|---|---|---|
| CaCTS | *ZBTB16, ZFHX3, BNC2, FOXO3, MBNL2, FOXP1, ZNF652, BHLHE41, FLI1, ZNF846, NCOA3, MXD4, RFX3, BAZ2B, ZEB1, MEF2C, MAF, BBX, ZBTB1, ZNF207, NCOA2, RUNX2, GATAD2B, FOXN3, ZNF148, ZNF609, SKI, BPTF, ZNF618, NFIA, FOXP2, ZBTB20, TET2, ZNF280D, CREB1, SPEN, ZNF518A, TET3, NRF1, SON, LCORL, ZNF33A, FOXJ3, AEBP2, REL, RFX7, TCF7L2, ARID2, SMYD3,* and *ZZZ3* | *ZBTB16, BNC2, MAF, FOXP1, FOXO3, ZNF846, ZFHX3, PPARG, FLI1, RUNX2, MBNL2, RFX3, MXD4, ZNF652, NFIA, JDP2, NCOA3, ADNP, ZNF710, MEF2C, SKIL, BBX, ATF6, BHLHE41,* and *TET2* |
| RegEnrich | *STAT1, STAT2, STAT3, BACH1, FOXP1, NFKB1, IRF1, ESR1, RUNX1, TAL1, FOS, ETS1, SP1, CEBPA, TP53, EGR1, E2F1, SPI1, CTCF, MITF, HIF1A, GATA2, RELA, FOXA1,* and *GATA3* | *STAT1, FOXP1, TAL1, GATA2, CEBPA, FOS, RUNX1, GATA3, CTCF, SPI1, FOXO3, MEF2C, ERG, STAT2, FOXA1, HNF4A, E2F4, FLI1, E2F1, NFYB, AR, BCL6, PPARG, TFAP2C,* and *BACH1* |
| PCSF | *ETV6, STAT1, PPARA, BACH1, STAT2,* and *MAFB* | *STAT1, MAFB, ETV6, JUNB, FOXO4,* and *STAT2* |
| TRIPS | *IRF1, HNF4A, CEBPA, RUNX1, STAT1, ESR1, HIF1A, PPARG, MYC, FOS, NFKB1, GATA3, STAT3, SPI1,* and *FOXP1* | *CEBPA, FOXP1, TP53, IRF1, STAT2, SPI1, FOXO3, HIF1A, NFKB1, FOXA1, ERG, TFAP2C, MYC, FOS, STAT1, HNF4A, TAL1,* and *ZNF550* |
| MOGAMUN | *FOS, ETS1, CUX1, FOXA1, FOXO3, FOXP1, GATA3, MYC, NFKB1, RUNX1, SMAD3, STAT3, TP53, CEBPA, E2F1, JUN, AR, ESR1, ETV6, SP1, MITF, BCL6, SPI1, IRF1, STAT1, HNF4A, TAL1, IRF9, STAT2,* and *GATA2* | *FOS, EPAS1, ETS1, CUX1, FOXP1, GATA3, MYC, NFKB1, RUNX1, SMAD3, STAT3, TP53, CEBPA, E2F1, JUN, HIF1A, RARA, AR, RELA, ESR1, ETV6, SP1, TFAP2C, SPI1, CTCF, IRF1, EGR1, STAT1, HNF4A, CREB1, MITF, TAL1, IRF9, STAT2, ZNF398,* and *E2F4* |

[S1] S. D. Ghiassian, J. Menche, and A.-L. Barabási. A DIseAse MOdule detection (DIAMOnD) algorithm derived from a systematic analysis of connectivity patterns of disease proteins in the human interactome. *PLoS Comput. Biol.*, 11(4): e1004120, Apr. 2015.
